# Clinical Response to Neurofeedback in Major Depression Relates to Subtypes of Whole-Brain Activation Patterns During Training

**DOI:** 10.1101/2024.05.01.592108

**Authors:** Masaya Misaki, Kymberly D. Young, Aki Tsuchiyagaito, Jonathan Savitz, Salvador M. Guinjoan

**Author notes:** Corresponding author: Masaya Misaki.

## Abstract

Major Depressive Disorder (MDD) poses a significant public health challenge due to its high prevalence and the substantial burden it places on individuals and healthcare systems. Real-time functional magnetic resonance imaging neurofeedback (rtfMRI-NF) shows promise as a treatment for this disorder, although its mechanisms of action remain unclear. This study investigated whole-brain response patterns during rtfMRI-NF training to explain interindividual variability in clinical efficacy in MDD. We analyzed data from 95 participants (67 active, 28 control) with MDD from previous rtfMRI-NF studies designed to increase left amygdala activation through positive autobiographical memory recall. Significant symptom reduction was observed in the active group (*t*=-4.404, *d*=-0.704, *p*<0.001) but not in the control group (*t*=-1.609, *d*=-0.430, *p*=0.111). However, left amygdala activation did not account for the variability in clinical efficacy. To elucidate the brain training process underlying the clinical effect, we examined whole-brain activation patterns during two critical phases of the neurofeedback procedure: activation during the self-regulation period, and transient responses to feedback signal presentations. Using a systematic process involving feature selection, manifold extraction, and clustering with cross-validation, we identified two subtypes of regulation activation and three subtypes of brain responses to feedback signals. These subtypes were significantly associated with the clinical effect (regulation subtype: *F*=8.735, *p*=0.005; feedback response subtype: *F*=5.326, *p*=0.008; subtypes’ interaction: *F*=3.471, *p*=0.039). Subtypes associated with significant symptom reduction were characterized by selective increases in control regions, including lateral prefrontal areas, and decreases in regions associated with self-referential thinking, such as default mode areas. These findings suggest that large-scale brain activity during training is more critical for clinical efficacy than the level of activation in the neurofeedback target region itself. Tailoring neurofeedback training to incorporate these patterns could significantly enhance its therapeutic efficacy.

## Introduction

Major Depressive Disorder (MDD) presents a significant public health challenge, with approximately one-third of diagnosed patients not responding to first-line treatments such as antidepressants and psychotherapy. This results in substantial disability and economic losses due to treatment costs and lost productivity ^1, 2^. Real-time functional magnetic resonance imaging neurofeedback (rtfMRI-NF) has emerged as a promising alternative, demonstrating large to medium effect sizes in treating depressive symptoms ^3-5^. This noninvasive brain modulation technique involves the real-time analysis and visualization of brain activation signals, thereby enabling participants to self-regulate their brain activity. Its efficacy in training participants to modulate their brain activation is well-supported by many studies, including several meta-analyses ^3-11^. However, the direct impact of rtfMRI-NF on symptom relief is not yet fully understood due to incomplete knowledge of the neural mechanisms by which this training alleviates symptoms through the regulation of specific brain activations.

Previous studies on the mechanisms of NF training ^12-15^, including investigations into brain responses to feedback signals ^15-19^, have identified a broad spectrum of brain activities associated with NF-mediated self-regulation training. This training process is considered to include aspects of reinforcement learning, and two types of brain activation - evaluation of feedback values and modulation of brain activation - are common components of the reinforcement learning process ^20^. Thus, to elucidate the learning mechanisms of NF-mediated brain regulation, investigating these two epochs is crucial. Active regions during these epochs typically include the prefrontal cortex, salience network, and reward processing areas ^12-15^. However, while many studies have focused on the success of regulating target brain activities, the relationship between these activities and subsequent symptom relief remains elusive. Furthermore, interindividual variability in the clinical efficacy of NF necessitates further investigation to identify brain response subtypes associated with therapeutic outcomes.

This study aimed to characterize whole-brain activation patterns during rtfMRI-NF training in individuals with MDD, with the goal of identifying brain activation subtypes associated with interindividual variability in therapeutic efficacy. To this end, we analyzed a large dataset from rtfMRI-NF studies where participants with MDD were trained to regulate left amygdala activity through neurofeedback ^21-24^. These studies consistently observed significant reductions in depressive symptoms on average post-training, albeit with variations in therapeutic outcomes among participants.

We hypothesized that the observed variability in treatment efficacy could be explained by variations in whole-brain activation patterns during self-regulation training, extending beyond the NF target region (amygdala). The involvement of large-scale networks in NF training has been demonstrated in a meta-analysis of NF studies ^12^, and burgeoning evidence suggests that the effects of NF training may extend beyond the targeted brain region ^25-27^. Specifically, we focused on two types of brain activation critical for NF training: regulatory task activity throughout the task block and instantaneous responses to neurofeedback signal presentations (Figure 1). These two types of activation are thought to correspond to two critical components of the reinforcement learning process ^20^ and should characterize the training process for each individual.

**Figure 1.**
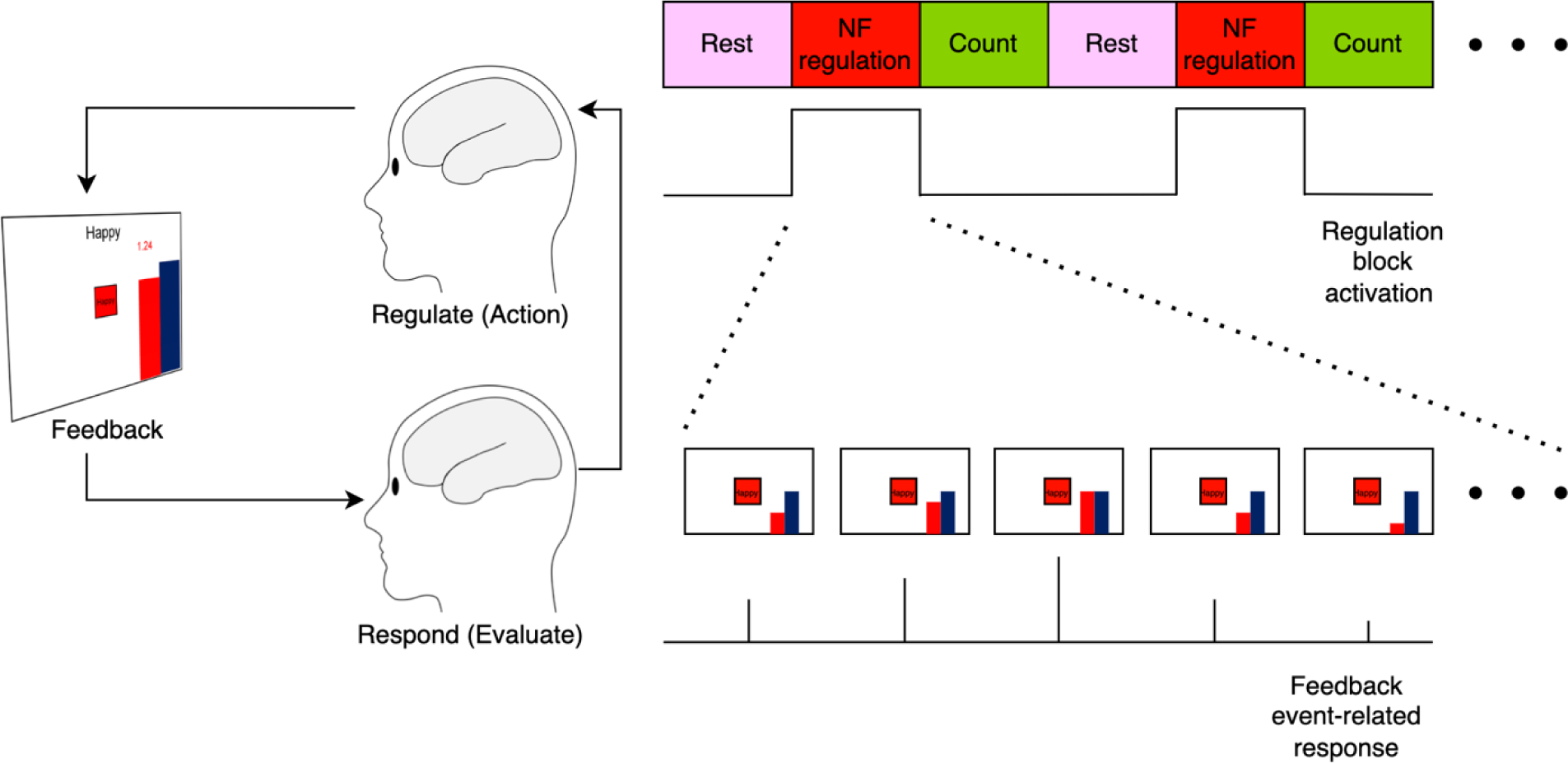
Schematic diagram of the neurofeedback-based brain regulation training loop (left panel) and the models used to estimate brain activation at each epoch (right panel). The reinforcement learning process is typically characterized by the evaluation of a feedback signal followed by the adjustment of action (mental regulation) to increase the reward (feedback) signal. Brain activations corresponding to these two components were analyzed using a block-wise response model during the neurofeedback regulation blocks and an event-related response model for each neurofeedback presentation at every TR, modulated by feedback amplitude.

## Methods

### Participants

The present study was a secondary analysis of data from our previously published studies ^21, 22, 24^ and a preliminary study utilizing the same rtfMRI-NF protocols in individuals diagnosed with MDD. The University of Oklahoma Institutional Review Board (IRB) or the Western IRB reviewed and approved the study protocols, ensuring adherence to the ethical principles of the Declaration of Helsinki. Participants provided written informed consent prior to participation and were financially compensated. Participants met DSM-IV-TR ^28^ criteria for MDD based on the Structural Clinical Interview for DSM-IV disorders ^29^ or DSM-5 criteria for MDD based on the Mini-International Neuropsychiatric Interview (MINI) ^30^. Previous articles ^21, 22, 24^ detailed each study’s inclusion and exclusion criteria. Common inclusion criteria across studies are ages 18-65, current diagnosis of MDD, and common exclusion criteria are current diagnosis of PTSD, substance use disorder, bipolar disorder, active suicidal ideation or behavior within a year, a history of psychosis, pregnancy, and MRI contraindicators.

The present analysis included data from 95 participants with MDD (68 females; mean age ± SD = 33.6 ± 10.4 years), consisting of 67 in the active NF group (46 females) who received left amygdala neurofeedback and 28 in the control group (22 females) who received neurofeedback from a brain region not associated with emotional processing.

### Real-time fMRI Neurofeedback Paradigm

The NF training was designed to enhance the activation signal in the left amygdala while recalling happy autobiographical memories ^31^. The task sequence utilized a blocked design consisting of alternating periods of rest, self-regulation with NF, and number counting, each lasting 40 seconds. This sequence was repeated four times in a single training run, with participants undergoing three such runs per session. Because the number of sessions differed among the studies, the present analyses were confined to data from the three runs of the initial session. Depressive symptom severity was estimated with the Montgomery-Åsberg Depression Rating Scale (MADRS) ^32^ immediately before the NF session, and one-week post-training.

MRI imaging in all experiments was conducted using the same 3T Discovery MR750 scanner (GE Healthcare). Blood Oxygenation Level-Dependent (BOLD) fMRI data acquisition employed a T2*-weighted gradient-echo planar imaging (EPI) sequence, with parameters set as follows: TR/TE = 2000/30 ms, acquisition matrix = 96 × 96, field of view (FOV)/slice thickness = 240 mm/2.9 mm, flip angle = 90°, and 34 axial slices, using a SENSE acceleration factor of 2. EPI images were then reconstructed to a 128x128 matrix, resulting in an fMRI voxel size of 1.875x1.875x2.9 mm³. Anatomical reference was obtained using a T1-weighted magnetization-prepared rapid gradient-echo (MPRAGE) sequence.

A detailed description of the rtfMRI-NF procedure can be found in our previous publications ^21, 22, 24^. Briefly, a custom in-house rtfMRI system was utilized for the experiments ^31^. For the active group, the NF signal was extracted from the left amygdala, defined by 7-mm diameter spheres centered at Talairach coordinates (x, y, z = -21, -5, -16 mm), and then mapped to each participant’s brain space. For the control group, the NF signal was sourced from the horizontal segment of the intraparietal sulcus at Talairach coordinates (-42, -48, 48 mm), a region suggested to be unrelated to emotion regulation ^33^. During the happy memory recall block, the NF signal was quantified as the percent signal change from the mean signal in the preceding rest block. The signal was updated every 2 s and visually presented to participants as a red bar.

### MRI data processing

The Analysis of Functional NeuroImages (AFNI; http://afni.nimh.nih.gov) software suite was utilized for image processing. After discarding the first three volumes to achieve signal equilibrium, preprocessing steps were conducted, including despike, RETROICOR ^34^ along with respiratory volume per time (RVT) regression ^35^ for physiological noise correction, slice-timing and motion corrections, nonlinear warping to the MNI template brain with resampling to 2 mm³ voxels using the Advanced Normalization Tools (ANTs; http://stnava.github.io/ANTs/), spatial smoothing with a 6 mm full-width at half maximum (FWHM) Gaussian kernel, and scaling of signals to percent change relative to the voxel-wise mean.

Activation during the self-regulation was assessed using a general linear model (GLM) analysis. We extracted the two types of brain response in this process (Fig. 1). The GLM regressors included response models for the regulation and counting blocks, each modeled with a boxcar function convolved with the canonical hemodynamic response function (HRF). The regressor for the regulation block was used for estimating the first type of brain activation, regulation block activation (Fig. 1). The GLM regressors also included twelve motion parameters (three rotations, three translations, and their temporal derivatives), three principal components of ventricle signals, local white matter signals (ANATICOR) ^36^, and an event-related regressor for the onset of any condition block (modeled as a delta function convolved with HRF) as nuisance covariates. Volumes with frame-wise displacement greater than 0.3 and their preceding volume were censored in the GLM.

Another type of brain response, the feedback event-related response (Figure 1), was evaluated using additional event-related regressors for each feedback presentation. The response was modeled as a delta function, modulated by the feedback amplitude normalized in each run, and convolved with the hemodynamic response function (HRF). Since the feedback signal was presented during the regulation block and this event regressor could be collinear with the regulation block regressor, we orthogonalized the feedback-response-event regressor with respect to the regulation block regressor. The beta coefficients from the general linear model (GLM) analysis were used as response estimates.

These response estimates were assessed for each block independently using the least squares - separate (LS-S) approach ^37^, in which separate regressors for the target block and all other blocks were included to estimate the response in a block, and repeated for each block in separate GLM analyses. The use of this block-wise response facilitates subsequent group analysis using linear mixed-effect model and provides the estimates with better test-retest reliability ^38, 39^.

### Clustering analysis

Clustering analysis was conducted on the whole-brain beta maps solely for the active group participants to categorize their brain activation patterns. This approach was chosen because including the control group could introduce brain activation patterns with various unspecific effects. Such inclusion might increase the dimensionality of latent subtypes and complicate the extraction of subtypes relevant to the active NF training process.

A significant challenge posed by this analysis is the high dimensionality of the whole-brain beta maps; the problem often referred to as the “curse of dimensionality” ^40^. This phenomenon refers to the issues arising in high-dimensional spaces, where distances between points become uniformly large, data points are sparsely distributed, and there is a high risk of overfitting, leading to clustering solutions that are difficult to reproduce.

To address this challenge, we implemented a multi-step strategy to reduce dimensionality and identify an informative space associated with treatment outcomes. Additionally, we employed cross-validation to ensure the clustering solution’s stability and reproducibility. Our approach involved (1) aggregating voxel-wise responses into regional averages using a functional brain atlas, (2) removing the regions irrelevant to treatment outcomes, (3) applying Uniform Manifold Approximation and Projection (UMAP) ^41^ to extract a low-dimensional representation, (4) applying k-means clustering in the UMAP space, and (5) employing repeated cross-validation to assess the robustness of the clustering solution ^42^. Figure 2 illustrates the flowchart of the analysis performed to delineate subtypes of brain activation.

**Figure 2.**
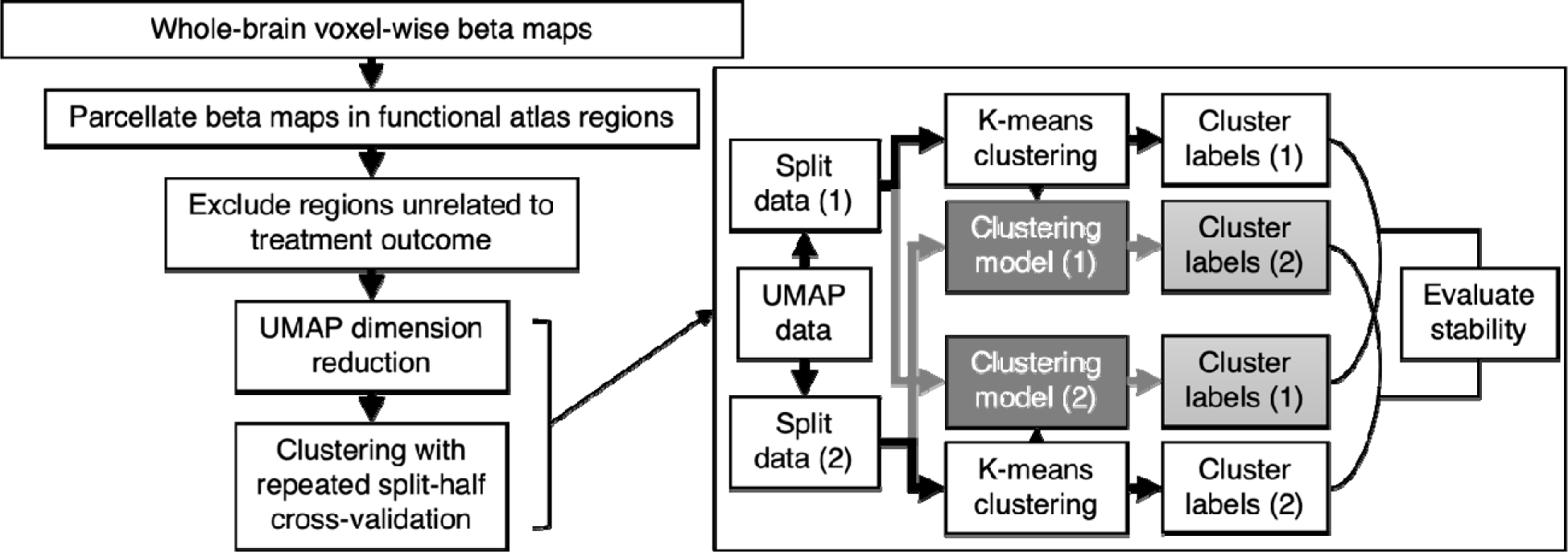
Procedures of subtyping whole-brain beta maps.

Initially, voxel-wise beta values were averaged within each functional region defined by the Shen 268 atlas ^43, 44^. This atlas was chosen for its demonstrated performance in various predictive modeling studies ^43, 45-49^. The data for each participant are averaged across the three runs or for each run. We tested both approaches as a hyperparameter to test which one yielded the most stable clustering solution. Subsequently, we identified regions correlating with changes in the MADRS score (change relative to the baseline score expressed as a ratio) after adjusting for the effects of age and sex. A threshold of *p* < 0.1 was used for this selection, prioritizing dimensionality reduction rather than finding significantly related regions. After excluding the regions irrelevant to the treatment outcome, we utilized UMAP for dimensionality reduction, followed by k-means clustering. The UMAP parameters were set to optimize the clustered distribution within the reduced-dimensional space, with ‘n_neighbors’ (the number of neighboring points used in the local manifold approximation) set to 50, and ‘min_dist’ (the minimum distance apart that points are allowed to be in the low-dimensional representation) set to 0. The large ‘n_neighbors’ value and small ‘min_dist’ value encourage the UMAP to extract a space with a clustered distribution. Subsequently, k-means clustering was executed within the UMAP-defined space.

The stability of the clusters was rigorously evaluated through cross-validation, facilitated by the ‘reval’ Python package ^42^. This involved dividing the dataset into two halves, applying clustering to one half, and then applying the derived clustering rule to the opposite half, and vice versa, to assess the consistency of the cluster labels between the two rules (Fig. 2). This procedure was repeated 10 times, each with a unique data split, to calculate the average stability score. The stability measure is the mean normalized Hamming distance between the two solutions, ranging from 0 to 1. A smaller value indicates a more stable and reproducible solution. Since this measure is larger with a larger number of clusters, it was scaled by the stability of random labeling of the same number of clusters ^50^.

In summary, these procedures were conducted across the hyperparameter space of response summary (average across runs or a sequence of three runs), UMAP random seed (50 values), UMAP dimension (ranging from 2 to 50), and the number of clusters (ranging from 2 to 5) to find robust clustering with the smallest mean distance between the cross-validated solutions.

### Mapping the brain responses in each subtype

After identifying the subtypes, we evaluated the voxel-wise response patterns for each subtype using a linear mixed-effect (LME) model analysis performed voxel-wise with the ‘lme4’ package ^51^ in R language and environment for statistical computing. This LME analysis was applied to the beta values for each block, run, and participant, incorporating fixed effects for the run, the identified subtype from the clustering analysis, age, and sex, as well as a random effect for participants at the intercept level. The mean response for each subtype was calculated using the ‘emmeans’ package ^52^ in R. The mean maps of the subtypes were thresholded at a voxel-wise *p* < 0.001, corrected for cluster size with a *p* < 0.05 using AFNI 3dClustSim.

### Statistical testing

LME analysis was performed to investigate changes in MADRS scores one week after a NF session for both active and control groups. The LME model included fixed effects for time (pre-/post-NF), group (active/control), age, and sex, as well as a random effect for participants at the intercept. We also examined whether treatment efficacy was correlated with NF training performance measures, including the mean NF amplitude, mean left amygdala response during the regulation block, and the change in left amygdala response between the first and the last training runs using linear model analysis.

We then examined whether the identified subtypes of brain activation during NF training were associated with demographic and training performance variables, as well as the depressive symptom score (MADRS) at the baseline and its change post-NF. Each variable was examined using linear model analysis, with subtypes serving as independent variables. The post-hoc analysis for the mean response for each subtype and the difference between the subtypes was performed using the ‘emmeans’ package ^52^, and *p*-values for the post-hoc comparisons were corrected by multivariate *t* adjustment.

In addition, the subtyping rules established in the active group were applied to the control group to assess if similar patterns in MADRS score changes could be observed across the subtypes. The presence of consistent subtype associations with symptom reduction in both the active and control groups would suggest the operation of a common treatment mechanism. This could indicate that the efficacy of NF training is not specific to the neurofeedback signal from an emotion-related region.

## Results

### Data selection

Post-training MADRS scores were not available for two participants in the active group. fMRI data from five participants were excluded from the analysis due to problems with physiological signal acquisition (two from the active group and one from the control group) or excessive head motion (two from the active group) with more than 30% of time points censored (Framewise Displacement [FD] > 0.3) in all training runs. Additionally, four participants in the active group who exhibited excessive head motion in any one of the training runs were excluded only when analyzing data with a series of three training runs. Analyses were performed using the largest sample size available for each test measure, regardless of missing data on other variables (see Supplementary Information [SI], Section 1). No significant differences in age, sex, and baseline MADRS scores were observed between the active and control groups in any of these selected datasets.

### Reduction of depressive symptoms following NF training

Figure 3 shows the MADRS scores before and after NF training for each group, with average scores represented by the height of bars and individual participant scores depicted as points connected by lines. LME analysis identified a significant main effect of time. Post-hoc analysis revealed a significant decrease in post-session scores (*t* = -4.404, *d* = -0.704, *p* < 0.001). SI Table S2a provides the ANOVA tables for the LME analysis. While the time by group interaction was not statistically significant (*p* = 0.090), the active group showed a significant symptom reduction (*t* = -5.576, *d* = -0.978, *p* < 0.001) but not the control group (*t* = -1.609, *d* = -0.430, *p* = 0.111).

**Figure 3:**
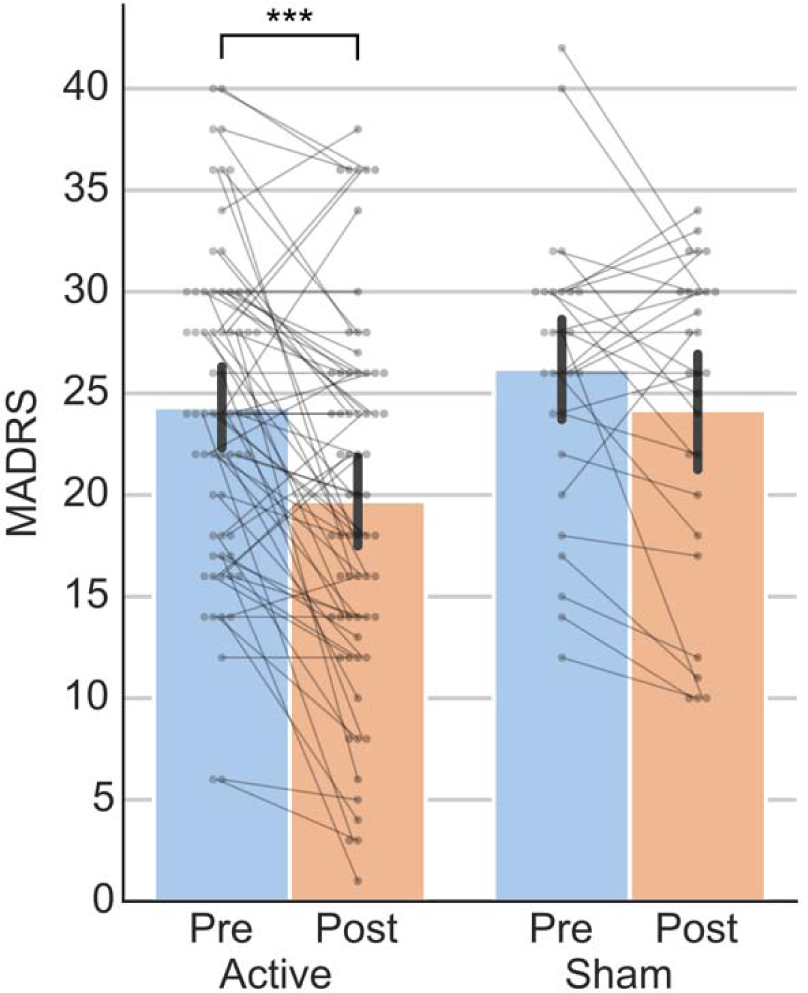
MADRS scores before and after NF training in the active and control groups. The bars show the mean MADRS scores for each group, with the error bars representing their standard error. Individual participant scores are shown as dots, with lines connecting pre- and post-training scores to highlight changes for each individual. A statistically significant decrease in post-training MADRS scores was observed in the active group (***, *p* < 0.001).

However, left amygdala activation had no significant main effect of the group (*F* = 2.870, *p* = 0.094) and the interaction between the group and run (*F* = 0.597, *p* = 0.551). Furthermore, within the active group, no significant associations were found between changes in MADRS scores and measures of NF training performance, including mean NF amplitude, average left amygdala response during regulation blocks, or changes in left amygdala response from the first to the last training runs (see SI Section 2 for comprehensive details).

### Clustering results

In clustering of activation maps related to the regulation block, the optimal stability score of 0.217 was obtained using the following hyperparameters: averaging responses across runs for each participant, extracting a 40-dimensional UMAP space, and clustering into two subtypes, which included 32 and 31 participants respectively. Similarly, in the clustering of activation maps associated with the feedback event, an optimal stability score of 0.229 was reached by analyzing a series of responses across three runs, extracting a 14-dimensional UMAP space, and forming three subtypes consisting of 23, 15, and 22 participants each. However, no significant association was found between the subtypes identified in the regulation block and those in the feedback event ( ^2^ = 3.143, *p* = 0.208).

Figure 4 shows *t*-maps of the mean beta values for the subtypes of regulation block activations (REG subtypes), with detailed peak coordinates available in SI, Section 3, Table S3. The first subtype, REG-A, exhibited increased BOLD signals in regions commonly identified in NF studies ^12^, including the lateral prefrontal cortex, superior and inferior parietal lobules, supplementary motor area (SMA), anterior insula, thalamus, and cerebellum. In contrast, decreased BOLD signals were observed in Heschl’s gyrus, posterior insula, middle cingulate cortex, and cuneus. The second subtype, REG-B, demonstrated increased BOLD signals in the lateral prefrontal cortex, SMA, and anterior insula, with decreased activity in default mode network (DMN) areas such as the precuneus, posterior cingulate cortex, mediodorsal prefrontal cortex, Rolandic operculum, and fusiform gyrus.

**Figure 4.**
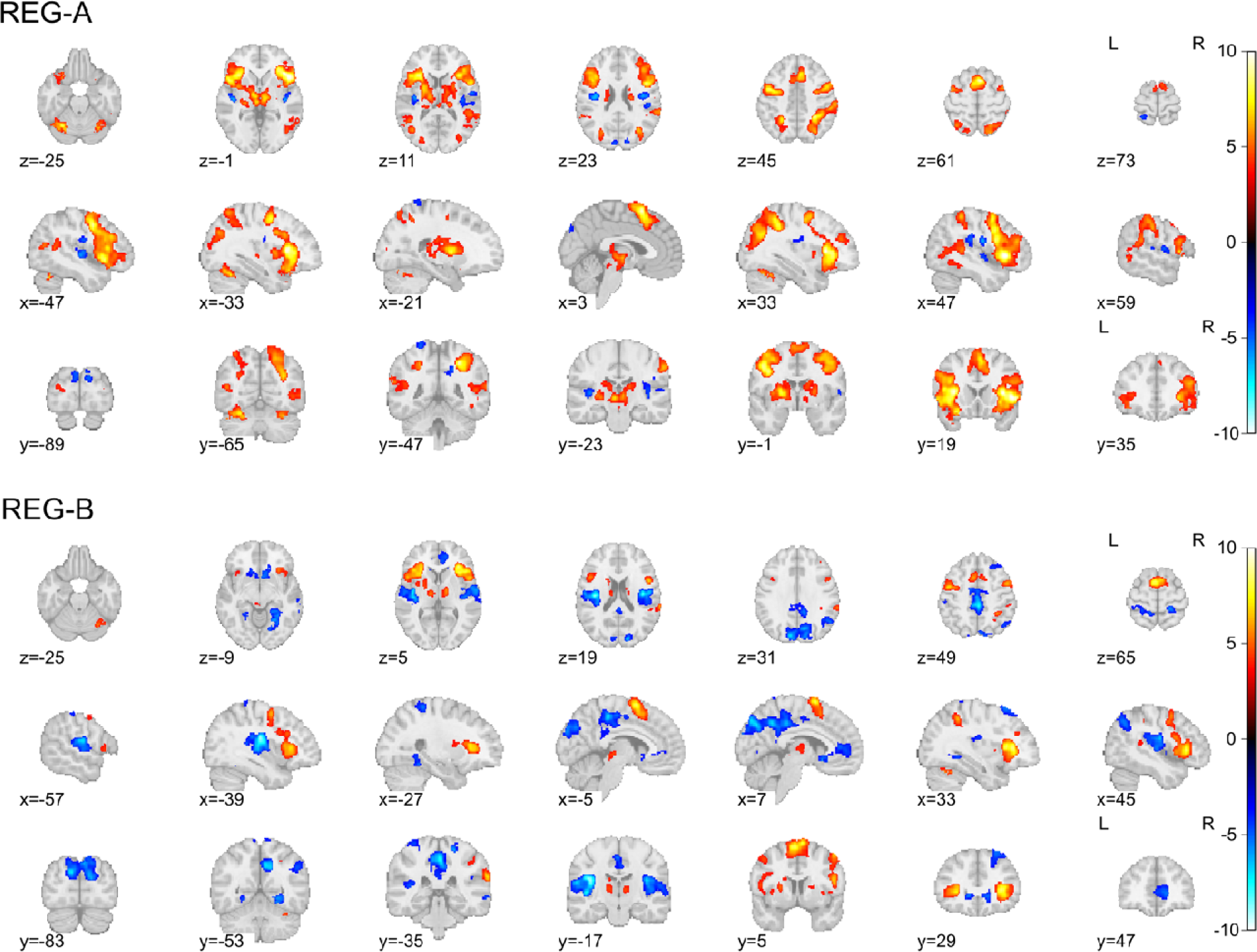
*t* maps of mean activation in the regulation block for the REG subtypes. The maps are thresholded at a voxel-wise p < 0.001, with cluster-size correction at p < 0.05.

Figure 5 presents *t*-maps of the mean beta values for subtypes of the brain responses to the feedback event (FBR subtypes), with peak coordinates detailed in SI, Section 4, Table S4. The first subtype, labeled FBR0, demonstrated a limited response, particularly in the Rolandic operculum and right supramarginal gyrus. The second subtype, labeled FBR-, was characterized by a negative association of BOLD signal changes in response to NF signals, affecting regions including the SMA, middle cingulate cortex, Heschl’s gyrus, Rolandic operculum, and visual cortex. In contrast, the third subtype, labeled FBR+, displayed a positive association of BOLD signal changes in areas such as the precuneus, middle cingulate cortex, lateral prefrontal cortex, nucleus accumbens, cerebellum, and inferior occipital regions.

**Figure 5.**
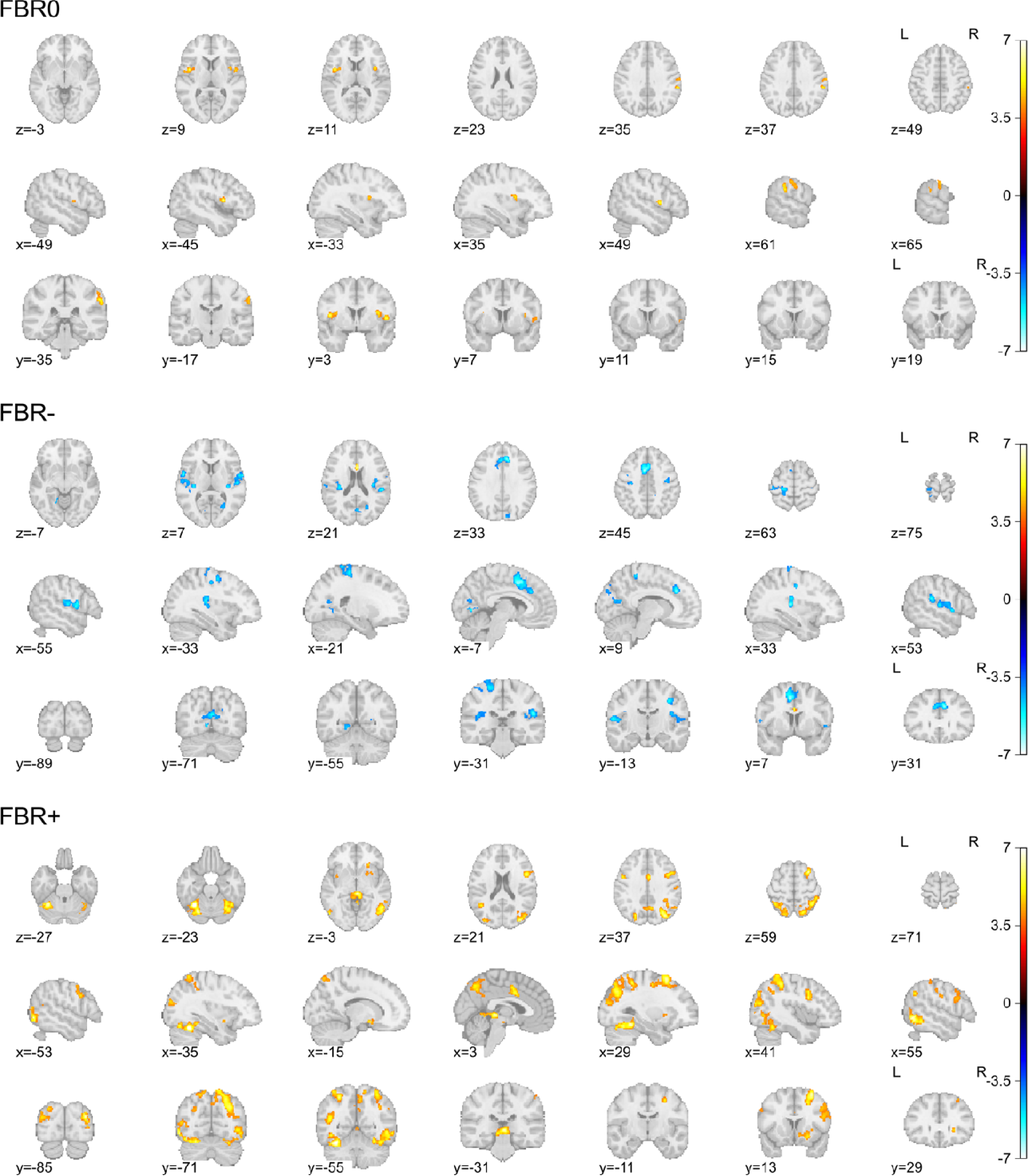
*t* maps of mean activation in response to NF signals for the FBR subtypes. The maps are thresholded at a voxel-wise *p* < 0.001, with cluster-size correction at *p* < 0.05.

### Subtype associations with demographics, NF-training characteristics, and the depressive symptom in the active group

Linear model analyses examining associations of the REG and FBR subtypes with age, mean NF signal amplitude, mean left amygdala activation across runs, and baseline MADRS scores revealed no significant differences between the subtypes. Similarly, ^2^ tests showed no significant associations of these subtypes with sex or study participation (refer to SI section 5 for further detailed statistics).

However, changes in MADRS scores, indicative of clinical efficacy, were significantly associated with these subtypes. The main effects of both REG and FBR subtypes, as well as their interaction effect on changes in MADRS scores, were statistically significant (REG, *F* = 8.735, *p* = 0.005; FBR, *F* = 5.326, *p* = 0.008; interaction *F* = 3.471, *p* = 0.039). Post-hoc analysis revealed a significant decrease in MADRS scores within the REG-B subtype (*t* = -6.077, *d* = -1.702, *p* < 0.001), in contrast to REG-A (*t* = -2.170, *d* = -0.608, *p* = 0.068). For the FBR subtypes, significant decreases in MADRS scores were observed in both FBR-(*t* = -4.995, *d* = - 1.399, *p* < 0.001) and FBR+ (*t* = -3.818, *d* = -1.069, *p* = 0.001), whereas the FBR0 subtype exhibited no significant change (*t* = -0.839, *d* = -0.235, *p* = 0.786).

Moreover, changes in MADRS scores varied by subtype combination, as illustrated in Figure 6. A significant reduction in scores was noted when REG-A was paired with FBR-(*t* = -3.327, *d* = -0.931, *p* = 0.003). Within the REG-B group, no significant reduction was observed with the FBR0 subtype (*t* = -1.221, *d* = -0.342, *p* = 0.401). However, significant reductions were seen when REG-B was paired with FBR-(*t* = -3.789, *d* = -1.061, *p* < 0.001) and FBR+ (*t* = -6.018, *d* = -1.685, *p* < 0.001). In summary, participants in the active group classified as REG-A with FBR-, or REG-B with FBR- or FBR+ experienced significant reductions in MADRS scores one week after NF training.

**Figure 6.**
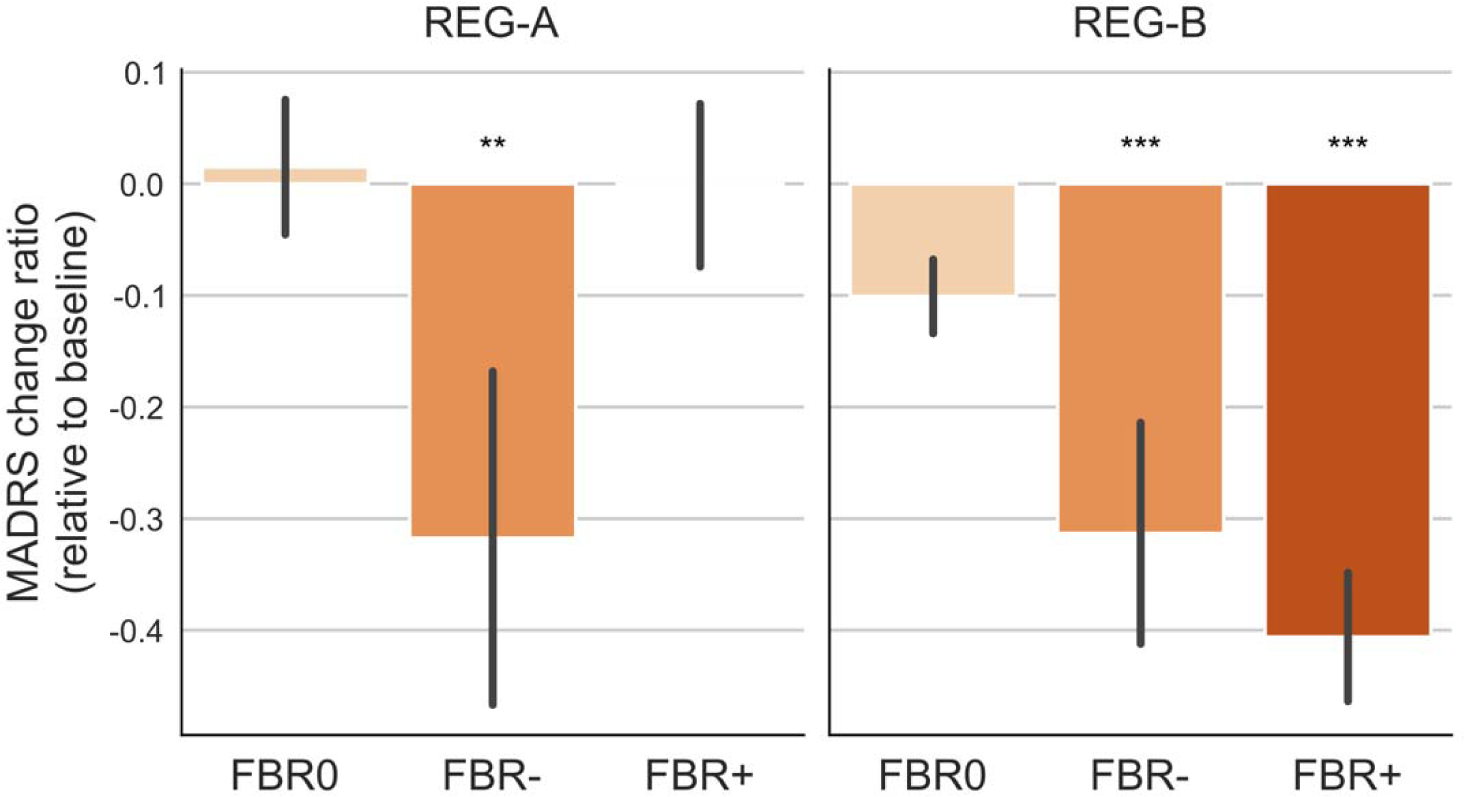
Ratio of change in MADRS score from baseline for each REG and FBR subtype. **, *p* < 0.005; ***, *p* < 0.001.

### Application of clustering rules to the control group

When the clustering rules derived from the active group were applied to the control group, participants were classified as follows: 13 into REG-A and 14 into REG-B for the REG subtypes; and 11 into FBR0, 5 into FBR-, and 11 into FBR+ for the FBR subtypes. No significant association was observed between the REG and FBR subtypes within the control group (χ² = 0.345, *p* = 0.842). Additionally, the distribution of participants across subtypes did not significantly differ from that observed in the active group for both REG (χ² = 0, *p* = 1.000) and FBR (χ² = 0.526, *p* = 0.768) subtypes.

In the control group, no significant associations were found between the subtypes and variables such as age, sex, study participation, mean NF signal amplitude, changes in left amygdala activation, or baseline MADRS scores, as detailed in SI Section 6. Contrary to the active group, there were no significant differences in MADRS score changes across subtypes within the control group.

## Discussion

The primary objective of this study was to identify subtypes of brain activation patterns during NF training that could explain interindividual differences in clinical response. Our analysis revealed that training characteristics within the target brain region, such as mean NF signal amplitude, mean left amygdala activation, and signal changes in the left amygdala, were not associated with changes in depressive symptoms. However, we found significant associations between subtypes of whole-brain activation patterns during NF training and changes in MADRS scores. In contrast, the control group showed no significant associations between the subtypes and changes in MADRS scores, underscoring that the effects of NF treatment are distinct from placebo effects.

Significant symptom reduction was observed across multiple subtype combinations, indicating the existence of multiple brain functional pathways to successful treatment. Notably, individuals classified within the FBR0 subtype, characterized by their non-responsiveness to feedback signals, did not experience significant symptom reduction, highlighting the critical role of brain response to feedback signals in effective NF training. Significant symptom reduction was evident in individuals exhibiting brain activation correlated with feedback signals, either in a positive (FBR+) or negative (FBR-) manner. However, the efficacy of this response pattern was dependent on their regulation subtype. Individuals exhibiting increased activation across broad brain areas during self-regulation (REG-A), which overlap with areas commonly reported in NF studies ^12^, required a negative response to NF signals (FBR-) to achieve significant symptom reduction. Conversely, individuals in the REG-B subtype, who showed increased activation in executive control areas such as the lateral prefrontal cortex and SMA and salience network regions such as the anterior insula (areas that overlap with those involved in general skill learning and emotion regulation ^53, 54^), alongside suppressive activation in DMN regions during self-regulation, achieved symptom reduction regardless of the polarity of their brain response to feedback signals. These findings suggest that nonspecific increases in brain activation do not contribute to treatment efficacy; rather, selective modulation of brain responses is crucial for successful NF therapy.

Increased activation of the DMN, often associated with self-referential thoughts ^55, 56^, has been frequently observed in individuals with MDD compared to healthy controls, and is linked to repetitive negative thinking ^57-65^. Particularly relevant to the pathophysiology of MDD and its treatment mechanisms could be the significant role of deactivating the posterior DMN, especially the posterior cingulate cortex (PCC). Current evidence suggests that the PCC acts as a major cortical hub for various self-referential phenomena, ranging from spatial body localization to self-talk and autobiographical information processing ^66-69^. In this context, the characteristic deactivation of the PCC observed in brain activation subtypes associated with clinical response in our study supports the hypothesis that this region may underlie the persistent and intensified negative self-referential mentation that characterizes depression.

The absence of suppressive activation during the regulation task can be compensated for by negative responses to feedback signals in the SMA, middle cingulate cortex (MCC), auditory cortex (including Heschl’s gyrus and Rolandic operculum), and visual cortex. Although the functional mechanism of these negative responses to NF signals remains unclear, they may counteract maladaptive brain activation patterns associated with MDD. Notably, alterations in SMA activity, frequently reported in MDD ^70-73^, and changes in middle cingulate activation during reward anticipation have also been highlighted in a meta-analysis ^74^. The association between less MCC activation during NF training and greater symptom reduction was also reported ^26^. Furthermore, increased connectivity in the auditory cortex was associated with repetitive negative thinking ^75^, and heightened activity or connectivity in the visual cortex have also been reported in individuals with MDD ^48, 76^.

Selective increases and decreases in brain activation associated with successful NF training have also been documented in a meta-analysis of amygdala NF studies ^4^. This study reported that successful amygdala modulation was linked to deactivation in the posterior insula and DMN regions, including the ventromedial prefrontal cortex and PCC, during the regulation period. Additionally, negative activations in successful modulators were noted in the left parahippocampal gyrus and cuneus, aligning with the present results observed in the FBR-subtype. Notably, this meta-analysis ^4^ defined training success as the ability to down-regulate amygdala activation, in contrast to our study, which focused on up-regulating amygdala activation and evaluated training success by changes in depressive symptoms. This suggests a common underlying mechanism in emotion regulation training that leads to depressive symptom reduction, despite varying approaches to NF signal definition and operational definitions of success.

In light of our findings, the therapeutic effects of NF targeting the left amygdala cannot be solely attributed to changes in its activity alone. This observation aligns with previous research showing that activations in brain regions beyond the intended target during NF training can mediate symptom reduction ^26^. Such evidence supports the notion that brain self-regulation through NF involves a large-scale network, extending well beyond a single target region. Although amygdala activity is a reliable marker of emotional states and can indicate successful regulatory efforts, as demonstrated in many NF protocols ^11^, our findings underscore the importance of the regulation process itself in achieving treatment success and explaining the observed variability in treatment outcomes.

To our knowledge, this study is the first to explore interindividual differences in brain activation during NF training and its association with therapeutic effects on depressive symptoms. This investigation marks a significant step toward understanding the mechanisms of NF as a treatment for psychiatric disorders. Our findings, which highlight the importance of the whole-brain regulation process over the mere amplitude of amygdala activity, could pave the way for improvements in NF protocols. Thus, incorporating feedback that targets both regulation-related activities and responses to feedback within large-scale brain networks could potentially enhance treatment efficacy. As NF methodologies continue to evolve, focusing on the process of regulating affective states rather than merely activating specific brain regions might yield more effective treatments. In this context, pattern-based NF approaches such as DecNef ^77^, connectome-based neurofeedback ^78^, and semantic neurofeedback ^79, 80^, which emphasize extracting feedback signals from multivariate brain activation patterns, offer a promising direction for refining training protocols.

The limitations of this study warrant careful consideration. Although the clustering analysis was designed to classify individuals into distinct subtypes, it is possible that these subtypes do not represent distinct groups per se. Rather, the subtypes may be part of a continuous distribution with no clear demarcation between groups. This is suggested by the fact that the obtained optimal stability scores, which indicate the discrepancy between cross-validated solutions, were greater than 0.2, indicating potential ambiguity in cluster definition. In scenarios where the samples have a continuous distribution, splitting into an equal number of participants could emerge as a more stable approach. This phenomenon may explain why our analysis resulted in nearly equal numbers of participants being categorized into each subtype, highlighting that these identified subtypes may not delineate distinct groups but rather capture different facets of the continuous response spectrum. It is also noteworthy that subtype classification was not related to age, sex, or baseline severity of depressive symptoms. However, the relationship between the subtypes and other demographic factors or neurobiological indicators before and after treatment remains unexplored. Further investigation may reveal subtypes of treatment response based on specific pretreatment characteristics.

In conclusion, the present study demonstrates that the therapeutic outcomes of NF training are significantly influenced by whole-brain activation patterns both during the process of affect regulation and during the response to feedback signals. Our findings reveal multiple patterns of brain activity associated with significant therapeutic effects, suggesting a variety of potential pathways to recovery through NF training. In future studies, the optimization of NF training may involve real-time monitoring of brain activity during regulation efforts and in response to feedback signals, as well as tailoring feedback signals to the individual’s current state of training progress. Such an approach could enhance treatment precision, adjusting it to match each participant’s unique neural response patterns, thus potentially increasing the effectiveness of NF training.

## Acknowledgement

This work was supported by the Laureate Institute for Brain Research, NIMH grant K99MH101235, R21MH113871, NIGMS grant P20GM121312, and NARSAD Young Investigator Grant from the Brain and Behavior Research Foundation. The funding sources had no role in the study design, data collection and analysis, decision to publish, or preparation of the manuscript. The authors are solely responsible for the content. The authors would like to acknowledge Jerzy Bodurka, Ph.D. (1964–2021), for his intellectual and scientific contributions to the designing the left amygdala neurofeedback protocols and development of the real-time fMRI neurofeedback systems, which provided the foundation for the present work. During the preparation of this work the authors used ChatGPT (https://chat.openai.com/) and DeepL (https://www.deepl.com/write) in order to improve language and readability. After using these tools, the authors reviewed and edited the content as needed and take full responsibility for the content of the publication.

