## Supplementary Information for "Clinical Response to Neurofeedback in Major Depression Relates to Subtypes of Whole-Brain Activation Patterns During Training"

### 1. Demographics for the selected datasets

Figure S1 shows the diagram illustrating the data selection process. Analyses were conducted using the maximum available sample size for each outcome measure, regardless of the presence of missing data on other variables. The analysis focusing on symptom change included all participants for whom post-training Montgomery-Åsberg Depression Rating Scale (MADRS) scores were available (n = 93). Clustering analyses of fMRI data included all eligible participants with available fMRI data, with two separate analyses: one for clustering the mean brain response across training runs (n = 63 in the active group and n = 27 in the control group) and another for clustering the sequence of brain responses across sessions (n = 57 in the control group and n = 27 in the control group). Combined analyses of both fMRI data and symptom change were conducted for participants with available post-training MADRS scores and fMRI data.

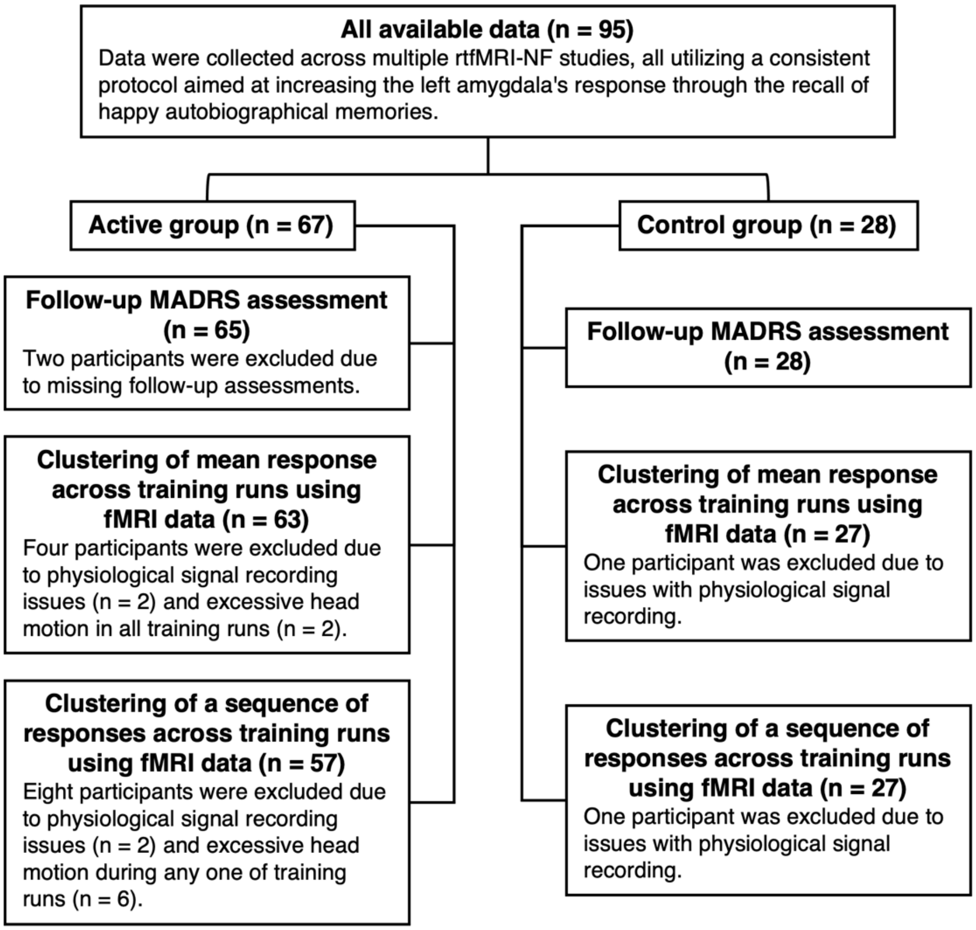

Figure S1. Diagram for the data selection.

No significant differences were observed between the active and control groups with respect to age, sex composition, and baseline MADRS scores. Table S1 details the comparative analyses. These variables showed no significant differences in any of the previously described data sets.

**Table S1.** Statistics of the comparative analyses for the dataset with post-training MADRS score.

| | Group | | Group difference |
|---|---|---|---|
| | **Active** | **Control** | **statistics** |
| N | 65 | 28 | |
| Age (mean ± SD) | 33.5 ± 10.4 | 33.9 ± 9.2 | $t$ = -0.148, $p$ = 0.883 |
| Sex (female:male) | 45:20 | 22:6 | $\chi^2$=0.848, $p$ = 0.357 |
| Baseline MADRS (mean ± SD) | 24.3 ± 7.7 | 26.2 ± 6.9 | $t$ = -1.164, $p$ = 0.249 |

### 2. MADRS change associations with amygdala self-regulation performance

The MADRS score decreased significantly after NF training. Table S2a shows the ANOVA table from the LME analysis for the MADRS score. The significant main effect of sex indicated that the MADRS scores were significantly higher for females than for males.

**Table S2a**. ANOVA table of the LME analysis for MADRS score.

| | SS | DF | *F* | *p* |
|---|---|---|---|---|
| Time | 434.863 | 1, 91 | 19.398 | < 0.001 |
| Group | 66.102 | 1, 90 | 2.949 | 0.089 |
| Age | 74.017 | 1, 89 | 3.302 | 0.073 |
| Sex | 132.642 | 1,89 | 5.917 | 0.017 |
| Time:Group | 65.895 | 1,91 | 2.939 | 0.090 |

SS, Sum of Squares; DF, Degree of Freedoms.

The change in MADRS scores (ratio relative to baseline) showed no association with measures of NF training performance, including mean NF amplitude (Table S2b), average left amygdala response during regulation blocks (Table S2c), and the change in left amygdala response from the first to the last training run (Table S2d).

**Table S2c**. ANOVA table of the linear model analysis for the MADRS change ratio with the mean NF amplitude.

| | SS | DF | *F* | *p* |
|---|---|---|---|---|
| Mean NF | 0.070 | 1, 78 | 0.951 | 0.332 |
| Group | 0.196 | 1, 78 | 2.675 | 0.106 |
| Age | 0.185 | 1, 78 | 2.521 | 0.116 |
| Sex | 0.055 | 1, 78 | 0.755 | 0.387 |
| Baseline MADRS | 0.003 | 1, 78 | 0.040 | 0.841 |
| Mean NF:Group | 0.001 | 1, 78 | 0.013 | 0.911 |

NF, neurofeedback signal.

**Table S2d**. ANOVA table of the linear model analysis for the MADRS change ratio with the mean left amygdala response during the regulation block.

| | SS | DF | *F* | *p* |
|---|---|---|---|---|
| Mean LA response | 0.015 | 1, 78 | 0.206 | 0.650 |
| Group | 0.155 | 1, 78 | 2.096 | 0.152 |
| Age | 0.162 | 1, 78 | 2.189 | 0.143 |
| Sex | 0.041 | 1, 78 | 0.559 | 0.457 |
| Baseline MADRS | 0.009 | 1, 78 | 0.125 | 0.725 |
| Mean NF:Group | 0.010 | 1, 78 | 0.134 | 0.715 |

LA, Left amygdala.

**Table S2e**. ANOVA table of the linear model analysis for the MADRS change ratio with the left amygdala response change between the first and the last training runs.

| | SS | DF | *F* | *p* |
|---|---|---|---|---|
| LA response change | 0.006 | 1, 78 | 0.078 | 0.780 |
| Group | 0.146 | 1, 78 | 1.979 | 0.163 |
| Age | 0.167 | 1, 78 | 2.267 | 0.136 |
| Sex | 0.030 | 1, 78 | 0.406 | 0.526 |
| Baseline MADRS | 0.007 | 1, 78 | 0.093 | 0.761 |
| Mean NF:Group | 0.021 | 1, 78 | 0.278 | 0.599 |

LA response change, Left amygdala response change between the first and the last training runs.

### 3. Peak coordinates of the statistical maps for the regulation subtypes

The peak coordinates in the mean beta maps for each regulation subtype are reported in tables S3a and S3b. Statistical maps were thresholded with voxel-wise $p < 0.001$ with cluster-size correction $p < 0.05$. The peaks with a minimum distance of 8 mm in each significant cluster are reported.

**Table S3a**. Peak coordinates of the mean beta map for the regulation subtype A

| Cluster ID | X | Y | Z | Peak Stat (*t*) | Cluster Size ($mm^3$) | Area |
|---|---|---|---|---|---|---|
| | Positive peaks | | | | | |
| 1 | 47 | 19 | -1 | 10.1245 | 138488 | Right Inferior Frontal Gyrus (p. Opercularis) |
| | 39 | 25 | -1 | 10.1101 | | Right Insula Lobe |
| | 33 | 21 | 7 | 9.9055 | | Right Insula Lobe |
| | -33 | 19 | 7 | 9.8553 | | Left Insula Lobe |
| 2 | 27 | -47 | 45 | 9.5005 | 42720 | Right Inferior Parietal Lobule |
| | 35 | -69 | 27 | 7.3018 | | Right Middle Occipital Gyrus |
| | 55 | -31 | 49 | 6.9776 | | Right Supra Marginal Gyrus |
| | 43 | -43 | 51 | 6.7953 | | Right Inferior Parietal Lobule |
| 3 | 3 | 13 | 61 | 9.4791 | 18584 | Right SMA |
| | 5 | 17 | 49 | 7.7881 | | Right SMA |
| | -7 | 9 | 59 | 7.6303 | | Left SMA |
| | -5 | -3 | 67 | 6.1768 | | Left SMA |
| 4 | -29 | -61 | -25 | 8.4624 | 5168 | Left Cerebellum (VI) |
| | -45 | -61 | -29 | 7.2203 | | Left Cerebellum (Crus 1) |
| | -25 | -71 | -23 | 6.6713 | | Left Cerebellum (VI) |
| | -41 | -69 | -17 | 4.8351 | | Left Fusiform Gyrus |
| 5 | -25 | -53 | 47 | 8.0507 | 12424 | Left Superior Parietal Lobule |
| | -31 | -75 | 31 | 6.4380 | | Left Middle Occipital Gyrus |
| | -35 | -61 | 59 | 6.1324 | | Left Superior Parietal Lobule |
| | -27 | -73 | 21 | 5.5020 | | Left Middle Occipital Gyrus |
| 6 | 31 | -65 | -25 | 6.6185 | 2536 | Right Cerebellum (VI) |
| | 33 | -53 | -29 | 6.1861 | | Right Cerebellum (VI) |
| | 27 | -65 | -25 | 5.9783 | | Right Cerebellum (VI) |
| | 43 | -57 | -29 | 5.5972 | | Right Cerebellum (Crus 1) |
| 7 | -45 | -67 | 13 | 5.9751 | 1432 | Left Middle Temporal Gyrus |
| 8 | -63 | -45 | 27 | 5.6263 | 3224 | Left Supra Marginal Gyrus |
| | -49 | -53 | 9 | 5.5665 | | Left Middle Temporal Gyrus |
| | -61 | -47 | 21 | 5.3385 | | Left Superior Temporal Gyrus |
| | -57 | -55 | 13 | 4.9804 | | Left Middle Temporal Gyrus |
| | Negative peaks | | | | | |
| 1 | -43 | -15 | 5 | -7.0113 | 4880 | Left Heschls Gyrus |
| | -39 | -15 | 19 | -6.6062 | | Left Rolandic Operculum |
| 2 | 41 | -19 | 1 | -6.7948 | 7744 | Right Insula Lobe |
| | 47 | -11 | 19 | -6.2492 | | Right Rolandic Operculum |
| | 39 | -11 | 21 | -6.1644 | | Right Insula Lobe |
| | 55 | -7 | 9 | -5.6869 | | Right Rolandic Operculum |
| 3 | 17 | -41 | 35 | -5.2424 | 960 | Right Middle Cingulate Cortex |

| | | | | | | |
|---|---|---|---|---|---|---|
| | 7 | -45 | 33 | -3.9672 | | Right Middle Cingulate Cortex |
| 4 | 11 | -89 | 27 | -5.2226 | 2632 | Right Cuneus |
| | -5 | -89 | 27 | -5.1304 | | Left Cuneus |
| | 11 | -89 | 33 | -4.9500 | | Right Cuneus |
| | 1 | -85 | 35 | -4.6507 | | Left Cuneus |
| 5 | -25 | -47 | 69 | -5.0963 | 704 | Left Superior Parietal Lobule |

SMA, Supplementary Motor Area.

**Table S3b**. Peak coordinates of the mean beta map for the regulation subtype B

| Cluster ID | X | Y | Z | Peak Stat (*t*) | Cluster Size ($mm^3$) | Area |
|---|---|---|---|---|---|---|
| | Positive peaks | | | | | |
| 1 | 33 | 29 | 5 | 8.628 | 20608 | Right Insula Lobe |
| | 45 | 23 | 3 | 8.492 | | Right Inferior Frontal Gyrus (p. Triangularis) |
| | 33 | 21 | 7 | 8.439 | | Right Insula Lobe |
| | 51 | 13 | 5 | 6.762 | | Right Inferior Frontal Gyrus (p. Opercularis) |
| 2 | -5 | 5 | 65 | 8.305 | 11936 | Left SMA |
| | 7 | 7 | 69 | 8.274 | | Right SMA |
| | 13 | 13 | 37 | 5.015 | | Right Middle Cingulate Cortex |
| 3 | -29 | 27 | 3 | 7.613 | 20328 | Left Insula Lobe |
| | -33 | 19 | 7 | 7.271 | | Left Insula Lobe |
| | -49 | 13 | 1 | 7.105 | | Left Inferior Frontal Gyrus (p. Opercularis) |
| | -41 | 23 | 3 | 7.073 | | Left Inferior Frontal Gyrus (p. Triangularis) |
| 4 | 67 | -35 | 23 | 7.062 | 2448 | Right Superior Temporal Gyrus |
| | 43 | -41 | 11 | 4.098 | | Right Superior Temporal Gyrus |
| 5 | -49 | -3 | 49 | 6.881 | 5528 | Left Precentral Gyrus |
| | -39 | -3 | 53 | 6.168 | | Left Precentral Gyrus |
| 6 | 31 | -45 | 45 | 6.446 | 2224 | Right Inferior Parietal Lobule |
| | 37 | -39 | 33 | 4.266 | | Right Angular Gyrus |
| | 41 | -37 | 41 | 4.171 | | Right Supra Marginal Gyrus |
| | 27 | -59 | 49 | 3.930 | | Right Superior Parietal Lobule |
| 7 | 13 | -5 | 9 | 6.190 | 3256 | Right Thalamus |
| | 23 | 9 | 11 | 4.496 | | Right Putamen |
| 8 | 31 | -63 | -27 | 5.825 | 1168 | Right Cerebellum (VI) |
| | 35 | -55 | -29 | 5.356 | | Right Cerebellum (VI) |
| | 39 | -63 | -25 | 4.747 | | Right Cerebellum (VI) |
| 9 | 37 | 39 | 27 | 4.788 | 584 | Right Middle Frontal Gyrus |
| 10 | -5 | -25 | -3 | 4.748 | 976 | Left Thalamus |
| | -11 | -31 | -15 | 3.991 | | Left Cerebellum (IV V) |
| | Negative peaks | | | | | |
| 1 | -41 | -17 | 21 | -9.181 | 15624 | Left Rolandic Operculum |
| | -43 | -13 | 7 | -7.950 | | Left Insula Lobe |
| | -57 | -15 | 15 | -6.974 | | Left Postcentral Gyrus |
| | -49 | -21 | 11 | -5.739 | | Left Superior Temporal Gyrus |
| 2 | 39 | -13 | 19 | -8.048 | 16032 | Right Insula Lobe |
| | 51 | -11 | 17 | -7.045 | | Right Rolandic Operculum |
| | 67 | -23 | -1 | -5.673 | | Right Superior Temporal Gyrus |
| | 39 | -21 | 1 | -5.613 | | Right Insula Lobe |
| 3 | 1 | -35 | 49 | -7.750 | 41104 | Left Middle Cingulate Cortex |
| | 9 | -73 | 37 | -7.224 | | Right Precuneus |
| | -11 | -83 | 29 | -7.199 | | Left Cuneus |
| | 13 | -51 | 37 | -7.153 | | Right Precuneus |

| | | | | | | |
|---|---|---|---|---|---|---|
| 4 | 21 | -37 | 61 | -7.102 | 2800 | Right Postcentral Gyrus |
| | 17 | -49 | 73 | -4.824 | | Right Postcentral Gyrus |
| 5 | 9 | 17 | -11 | -6.203 | 9144 | Right Olfactory cortex |
| | -13 | 27 | -3 | -5.979 | | Left Caudate Nucleus |
| | 13 | 47 | 1 | -5.445 | | Right Superior Medial Gyrus |
| | 5 | 39 | 7 | -5.258 | | Right Anterior Cingulate Cortex |
| 6 | 23 | -55 | -7 | -6.136 | 5544 | Right Lingual Gyrus |
| | 15 | -73 | -5 | -5.946 | | Right Lingual Gyrus |
| | 29 | -51 | -9 | -5.791 | | Right Fusiform Gyrus |
| | 21 | -49 | -11 | -5.749 | | Right Fusiform Gyrus |
| 7 | 31 | 21 | 59 | -5.996 | 3352 | Right Middle Frontal Gyrus |
| | 35 | 31 | 53 | -4.660 | | Right Middle Frontal Gyrus |
| | 21 | 29 | 41 | -4.283 | | Right Superior Frontal Gyrus |
| 8 | 43 | -63 | 39 | -5.612 | 5216 | Right Angular Gyrus |
| | 51 | -61 | 41 | -5.493 | | Right Angular Gyrus |
| | 53 | -61 | 29 | -5.365 | | Right Angular Gyrus |
| 9 | -25 | -39 | 61 | -5.538 | 3472 | Left Postcentral Gyrus |
| | -13 | -43 | 79 | -4.989 | | Left Postcentral Gyrus |
| | -13 | -45 | 65 | -4.975 | | Left Precuneus |
| | -33 | -33 | 69 | -4.862 | | Left Postcentral Gyrus |
| 10 | -27 | -45 | -17 | -5.179 | 1448 | Left Fusiform Gyrus |
| | -25 | -49 | -5 | -5.088 | | Left Lingual Gyrus |
| | -15 | -47 | -7 | -3.930 | | Left Lingual Gyrus |
| 11 | -53 | -27 | 53 | -4.981 | 696 | Left Postcentral Gyrus |

### 4. Peak coordinates of the statistical maps for the NF-response subtypes

The peak coordinates in the mean beta maps for each NF-response subtype are reported in tables S4a, S4b, and S4c. Statistical maps were thresholded with voxel-wise $p < 0.001$ with cluster-size correction $p < 0.05$. The peaks with a minimum distance of 8 mm in each significant cluster are reported.

**Table S4a**. Peak coordinates of the mean beta map for the NF-response subtype A

| **Cluster ID** | **X** | **Y** | **Z** | **Peak Stat (*t*)** | **Cluster Size (mm$^3$)** | **Area** |
|---|---|---|---|---|---|---|
| | Positive peaks | | | | | |
| 1 | -45 | 3 | 11 | 5.835 | 984 | Left Rolandic Operculum |
| | -53 | -1 | 7 | 4.659 | | Left Rolandic Operculum |
| | -33 | 3 | 15 | 4.593 | | Left Insula Lobe |
| 2 | 49 | 1 | 7 | 5.710 | 776 | Right Rolandic Operculum |
| | 53 | 9 | 1 | 4.537 | | Right Rolandic Operculum |
| 3 | 59 | -35 | 37 | 5.463 | 2048 | Right Supra Marginal Gyrus |
| | 63 | -33 | 27 | 5.213 | | Right Supra Marginal Gyrus |
| | 55 | -31 | 41 | 5.170 | | Right Supra Marginal Gyrus |
| | 63 | -19 | 39 | 4.727 | | Right Postcentral Gyrus |
| 4 | 37 | 5 | 11 | 5.254 | 832 | Right Insula Lobe |
| | 39 | -3 | 19 | 5.072 | | Right Rolandic Operculum |

**Table S4b**. Peak coordinates of the mean beta map for the NF-response subtype B

| **Cluster ID** | **X** | **Y** | **Z** | **Peak Stat (*t*)** | **Cluster Size (mm$^3$)** | **Area** |
|---|---|---|---|---|---|---|
| | Positive peaks | | | | | |
| 1 | 1 | 15 | 21 | 6.470 | 1232 | Right Anterior Cingulate Cortex |
| | 1 | 3 | 27 | 5.735 | | Left Anterior Cingulate Cortex |

| | | | | | | |
|---|---|---|---|---|---|---|
| | Negative peaks | | | | | |
| 1 | -7 | 7 | 45 | -6.363 | 8632 | Left SMA |
| | 1 | 15 | 47 | -6.264 | | Left SMA |
| | 9 | 31 | 31 | -6.078 | | Right Middle Cingulate Cortex |
| | -9 | 13 | 39 | -5.377 | | Left Middle Cingulate Cortex |
| 2 | 33 | -25 | 9 | -5.809 | 7472 | Right Insula Lobe |
| | 53 | -31 | 19 | -5.766 | | Right Superior Temporal Gyrus |
| | 63 | -3 | 7 | -5.454 | | Right Heschls Gyrus |
| | 53 | -17 | 7 | -5.101 | | Right Superior Temporal Gyrus |
| 3 | -55 | 1 | 7 | -5.607 | 6896 | Left Rolandic Operculum |
| | -59 | -13 | 11 | -5.434 | | Left Superior Temporal Gyrus |
| | -63 | -21 | 17 | -5.428 | | Left Postcentral Gyrus |
| | -33 | -27 | 15 | -5.366 | | Left Heschls Gyrus |
| 4 | 37 | -15 | 41 | -5.445 | 992 | Right Precentral Gyrus |
| 5 | 3 | -71 | 17 | -5.319 | 4192 | Right Calcarine Gyrus |
| | 17 | -67 | 15 | -5.170 | | Right Calcarine Gyrus |
| | 25 | -65 | 7 | -4.539 | | Right Calcarine Gyrus |
| | -17 | -65 | 11 | -4.508 | | Left Calcarine Gyrus |
| 6 | -31 | -7 | 53 | -5.308 | 1224 | Left Precentral Gyrus |
| | -37 | -3 | 49 | -4.353 | | Left Precentral Gyrus |
| 7 | -7 | -71 | -1 | -5.282 | 1280 | Left Lingual Gyrus |
| | -15 | -53 | -5 | -4.900 | | Left Lingual Gyrus |
| | -9 | -61 | 1 | -4.493 | | Left Lingual Gyrus |
| 8 | -21 | -41 | 73 | -5.224 | 3328 | Left Postcentral Gyrus |
| | -23 | -29 | 65 | -5.153 | | Left Postcentral Gyrus |
| | -43 | -27 | 61 | -5.101 | | Left Postcentral Gyrus |
| | -23 | -37 | 59 | -4.517 | | Left Postcentral Gyrus |
| 9 | 15 | -45 | 49 | -5.043 | 848 | Right Precuneus |
| | 9 | -39 | 57 | -4.928 | | Right Paracentral Lobule |
| 10 | -33 | -19 | 47 | -4.826 | 624 | Left Precentral Gyrus |
| | -41 | -21 | 37 | -4.492 | | Left Postcentral Gyrus |
| 11 | 13 | -81 | 37 | -4.695 | 728 | Right Cuneus |
| | 7 | -85 | 31 | -4.485 | | Right Cuneus |
| 12 | 35 | -27 | 67 | -4.611 | 576 | Right Precentral Gyrus |
| | 39 | -19 | 67 | -4.028 | | Right Precentral Gyrus |

**Table S4c**. Peak coordinates of the mean beta map for the NF-response subtype C

| Cluster ID | X | Y | Z | Peak Stat (t) | Cluster Size ($mm^3$) | Area |
|---|---|---|---|---|---|---|
| | Positive peaks | | | | | |
| 1 | -35 | -55 | -23 | 7.461 | 15720 | Left Cerebellum (VI) |
| | -27 | -69 | -21 | 6.839 | | Left Cerebellum (VI) |
| | -53 | -71 | -7 | 6.493 | | Left Inferior Occipital Gyrus |
| | -41 | -67 | -15 | 6.075 | | Left Fusiform Gyrus |
| 2 | 29 | -47 | -19 | 6.871 | 13032 | Right Fusiform Gyrus |
| | 53 | -63 | -9 | 6.840 | | Right Inferior Temporal Gyrus |
| | 31 | -59 | -21 | 6.241 | | Right Cerebellum (VI) |
| | 47 | -61 | -7 | 5.926 | | Right Inferior Temporal Gyrus |
| 3 | 37 | -85 | 21 | 6.722 | 28264 | Right Middle Occipital Gyrus |
| | 33 | -51 | 65 | 6.524 | | Right Superior Parietal Lobule |
| | 35 | -77 | 37 | 6.406 | | Right Middle Occipital Gyrus |
| | 37 | -51 | 59 | 6.256 | | Right Superior Parietal Lobule |
| 4 | 3 | -31 | -3 | 6.582 | 3808 | Cerebellar Vermis (3) |
| | 1 | -53 | -1 | 5.439 | | Cerebellar Vermis (4/5) |
| | -9 | -31 | -9 | 5.367 | | Left Lingual Gyrus |
| | 1 | -49 | -1 | 5.080 | | Cerebellar Vermis (4/5) |
| 5 | 29 | 13 | 59 | 6.369 | 5080 | Right Superior Frontal Gyrus |
| | 23 | 11 | 43 | 5.805 | | Right Superior Frontal Gyrus |
| | 29 | -11 | 49 | 5.158 | | Right Precentral Gyrus |
| | 27 | -1 | 49 | 4.487 | | Right Middle Frontal Gyrus |
| 6 | -29 | -53 | 53 | 6.291 | 7248 | Left Inferior Parietal Lobule |
| | -37 | -53 | 61 | 5.536 | | Left Superior Parietal Lobule |
| | -25 | -75 | 37 | 5.432 | | Left Superior Occipital Gyrus |
| | -23 | -79 | 51 | 5.307 | | Left Superior Parietal Lobule |
| 7 | -15 | 5 | -13 | 5.899 | 1784 | Left Olfactory cortex |
| | -29 | 3 | -11 | 5.402 | | Left Putamen |
| | -17 | 7 | -17 | 4.675 | | Left Olfactory cortex |
| | -41 | 5 | -5 | 4.578 | | Left Insula Lobe |
| 8 | 1 | 5 | 33 | 5.827 | 1608 | Left Middle Cingulate Cortex |
| | 3 | -1 | 45 | 5.290 | | Right Middle Cingulate Cortex |
| 9 | 41 | 5 | 27 | 5.798 | 5272 | Right Inferior Frontal Gyrus (p. Opercularis) |
| | 49 | 7 | 21 | 5.773 | | Right Inferior Frontal Gyrus (p. Opercularis) |
| | 43 | 7 | 35 | 5.559 | | Right Precentral Gyrus |
| | 59 | 13 | 21 | 4.218 | | Right Inferior Frontal Gyrus (p. Opercularis) |
| 10 | 21 | 7 | -13 | 5.787 | 4344 | Right Olfactory cortex |
| | 13 | -1 | -15 | 5.510 | | Right Hippocampus |
| | 23 | 15 | -13 | 5.431 | | Right Rectal Gyrus |

| | | | | | | |
|---|---|---|---|---|---|---|
| | 13 | -5 | -13 | 5.227 | | Right Hippocampus |
| 11 | -49 | 1 | 43 | 5.425 | 1576 | Left Precentral Gyrus |
| | -51 | 3 | 35 | 5.316 | | Left Precentral Gyrus |
| | -55 | 9 | 31 | 4.984 | | Left Precentral Gyrus |
| 12 | -7 | -17 | -15 | 4.828 | 592 | Left Ventral DC |
| | 7 | -15 | -13 | 4.564 | | Right Ventral DC |
| | -3 | -23 | -15 | 4.476 | | Brain Stem |
| 13 | 57 | -15 | 31 | 4.743 | 904 | Right Postcentral Gyrus |
| | 61 | -21 | 41 | 4.601 | | Right Supra Marginal Gyrus |
| | 63 | -17 | 27 | 4.221 | | Right Supra Marginal Gyrus |

### 5. Regulation and NF-response subtypes' associations with demographics and NF training characteristics

The subtypes showed no significant association with sex ($\chi^2 < 0.001$, p = 1.000 for the regulation subtype and $\chi^2 = 3.598$, p = 0.166 for the NF-response subtype) and the study difference ($\chi^2 = 1.651$, p = 0.648 for the regulation subtype and $\chi^2 = 1.876$, p = 0.931 for the NF-response subtype). Linear model analyses testing the association of these subtypes with age, mean NF signal amplitude, mean left amygdala activation, and the baseline MADRS score showed no significant difference between the subtypes (See tables S5a to S5d for ANOVA tables).

The difference in the left amygdala activation changes (3rd – 1st run) were observed between the NF-response subtypes (Table S5e). The NF-response subtype A showed left amygdala signal decrease while subtype C showed signal increase between the runs (Figure S2). However, the change in each subtype was not significant (A, $t$ = -1.507, $d$ = -0.414, $p$ = 0.355; B, $t$ = -0.008, $d$ = -0.037, $p$ = 0.999; C, $t$ = -2.299, $d$ = 0.631, $p$ = 0.074).

**Table S5a**. ANOVA table for age.

| | **SS** | **DF** | ***F*** | ***p*** |
|---|---|---|---|---|
| Regulation Subtype | 46.214 | 1,53 | 0.439 | 0.511 |
| NF-response subtype | 318.586 | 2,53 | 1.513 | 0.230 |
| Regulation Subtype x NF-response subtype | 159.178 | 2,53 | 0.756 | 0.475 |

SS, Sum of squares; DF, Degree of Freedoms

**Table S5b**. ANOVA table for the mean NF signal amplitude.

| | **SS** | **DF** | ***F*** | ***p*** |
|---|---|---|---|---|
| Regulation Subtype | 0.0430 | 1,53 | 0.3972 | 0.5312 |
| NF-response subtype | 0.1935 | 2,53 | 0.8941 | 0.4150 |
| Regulation Subtype x NF-response subtype | 0.2476 | 2,53 | 1.1440 | 0.3262 |

**Table S5c**. ANOVA table for the mean left amygdala activation during the regulation block.

| | **SS** | **DF** | ***F*** | ***p*** |
|---|---|---|---|---|
| Regulation Subtype | 0.0101 | 1,53 | 0.2121 | 0.6469 |
| NF-response subtype | 0.2084 | 2,53 | 2.1702 | 0.1241 |
| Regulation Subtype x NF-response subtype | 0.1654 | 2,53 | 1.7225 | 0.1884 |

**Table S5d**. ANOVA table for the baseline MADRS score.

| | **SS** | **DF** | ***F*** | ***p*** |
|---|---|---|---|---|
| Regulation Subtype | 3.396 | 1,53 | 0.057 | 0.812 |
| NF-response subtype | 45.919 | 2,53 | 0.388 | 0.681 |
| Regulation Subtype x NF-response subtype | 109.411 | 2,53 | 0.924 | 0.403 |

**Table S5e**. ANOVA table for the left amygdala activation change (3rd run – 1st run)

| | SS | DF | *F* | *p* |
|---|---|---|---|---|
| Regulation Subtype | 0.2152 | 1,53 | 3.8121 | 0.0561 |
| NF-response subtype | 0.4393 | 2,53 | 3.8911 | 0.0264* |
| Regulation Subtype x NF-response subtype | 0.1605 | 2,53 | 1.4215 | 0.2503 |

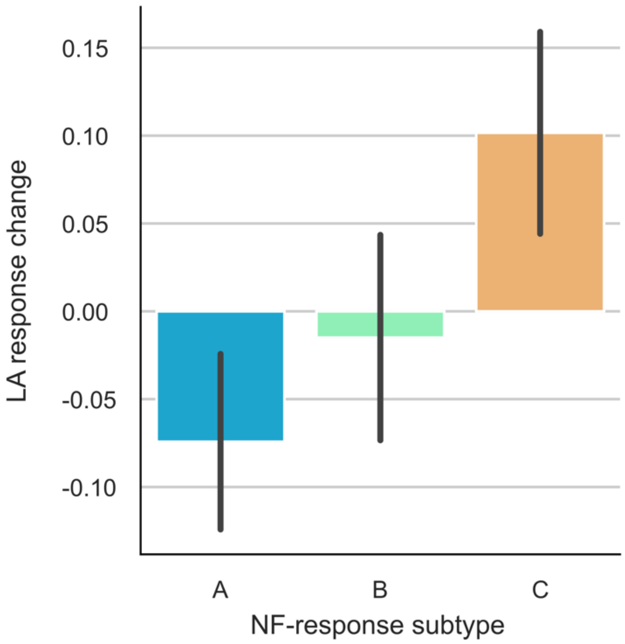

**Figure S2**. Left amygdala activation changes for each NF-response subtype.

### 6. Subtypes in the control control group

The subtypes in the control group showed no significant association with sex ($\chi^2$ = 2.228, *p* = 0.136 for the regulation subtype and $\chi^2$ = 2.384, *p* = 0.304 for the NF-response subtype) and the study difference ($\chi^2$ = 0.046, *p* = 0.830 for the regulation subtype and $\chi^2$ = 3.142, *p* = 0.208 for the NF-response subtype). Linear model analyses testing the association of these subtypes with age, mean NF signal amplitude, left amygdala activation change, the baseline MADRS score, and MDRS score change showed no significant difference between the subtypes (See tables S6a to S6f for ANOVA tables). Only a significant effect was found for the mean left amygdala activation during the self-regulation block in the interaction between the regulation and NF-response subtypes (Table S6c). Post-hoc analysis indicated that within the regulation subtype B, NF-response subtype B has larger left amygdala activation than subtype C (*t* = 2.558, *d* = 2.089, *p* = 0.045).

**Table S6a**. ANOVA table for age.

| | SS | DF | *F* | *p* |
|---|---|---|---|---|
| Regulation Subtype | 18.183 | 1,21 | 0.224 | 0.641 |
| NF-response subtype | 210.426 | 2,21 | 1.297 | 0.294 |
| Regulation Subtype x NF-response subtype | 25.638 | 2,21 | 0.158 | 0.855 |

SS, Sum of squares; DF, Degree of Freedoms

**Table S6b**. ANOVA table for the mean NF signal amplitude.

| | SS | DF | *F* | *p* |
|---|---|---|---|---|
| Regulation Subtype | 0.0977 | 1,21 | 2.6417 | 0.1190 |
| NF-response subtype | 0.0963 | 2,21 | 1.3007 | 0.2933 |
| Regulation Subtype x NF-response subtype | 0.0110 | 2,21 | 0.1497 | 0.8618 |

**Table S6c**. ANOVA table for the mean left amygdala activation during the regulation block.

| | SS | DF | *F* | *p* |
|---|---|---|---|---|
| Regulation Subtype | 0.0121 | 1,21 | 0.9092 | 0.3511 |
| NF-response subtype | 0.0097 | 2,21 | 0.3642 | 0.6990 |
| Regulation Subtype x NF-response subtype | 0.1084 | 2,21 | 4.0677 | 0.0321* |

**Table S6d**. ANOVA table for the left amygdala activation change (3rd run – 1st run)

| | SS | DF | *F* | *p* |
|---|---|---|---|---|
| Regulation Subtype | 0.1383 | 1,21 | 1.8072 | 0.1931 |
| NF-response subtype | 0.2798 | 2,21 | 1.8281 | 0.1853 |
| Regulation Subtype x NF-response subtype | 0.2789 | 2,21 | 1.8227 | 0.1862 |

**Table S6e**. ANOVA table for the baseline MADRS score.

| | SS | DF | *F* | *p* |
|---|---|---|---|---|
| Regulation Subtype | 14.2720 | 1,21 | 0.2402 | 0.6290 |
| NF-response subtype | 11.4716 | 2,21 | 0.0965 | 0.9083 |
| Regulation Subtype x NF-response subtype | 30.1400 | 2,21 | 0.2537 | 0.7782 |

**Table S6f**. ANOVA table for the MADRS score change ratio relative to baseline.

| | SS | DF | *F* | *p* |
|---|---|---|---|---|
| Regulation Subtype | 0.0060 | 1,21 | 0.1146 | 0.7383 |
| NF-response subtype | 0.0046 | 2,21 | 0.0441 | 0.9569 |
| Regulation Subtype x NF-response subtype | 0.0033 | 2,21 | 0.0320 | 0.9684 |
